# Mouse Academy: high-throughput automated training and trial-by-trial behavioral analysis during learning

**DOI:** 10.1101/467878

**Authors:** Mu Qiao, Tony Zhang, Cristina Segalin, Sarah Sam, Pietro Perona, Markus Meister

## Abstract

Progress in understanding how individual animals learn will require high-throughput standardized methods for behavioral training but also advances in the analysis of the resulting behavioral data. In the course of training with multiple trials, an animal may change its behavior abruptly, and capturing such events calls for a trial-by-trial analysis of the animal’s strategy. To address this challenge, we developed an integrated platform for automated animal training and analysis of behavioral data. A low-cost and space-efficient apparatus serves to train entire cohorts of mice on a decision-making task under identical conditions. A generalized linear model (GLM) analyzes each animal’s performance at single-trial resolution. This model infers the momentary decision-making strategy and can predict the animal’s choice on each trial with an accuracy of ~80%. We also assess the animal’s detailed trajectories and body poses within the apparatus. Unsupervised analysis of these features revealed unusual trajectories that represent hesitation in the response. This integrated hardware/software platform promises to accelerate the understanding of animal learning.

## INTRODUCTION

Learning – the change of neural representation and behavior that results from past experience and the consequences of actions – is important for animals to survive and forms a central topic in neuroscience^1^. Different individuals may apply different strategies to the learning process, reflecting their individual personalities. Indeed, substantial differences in sensory biases, locomotion, motivation, and cognitive competence have been observed in populations of fruit flies^2,3^, rodents and primates^4–6^. Thus, it is critical to investigate learning at the individual level.

Rodents, especially the mouse, have become popular experimental animals in studying associative learning and decision-making, because of the wide availability of transgenic resources^7–10^. They can learn to perform complex decision-making tasks that probe cognitive components such as working memory and selective attention^11–13^. However, differences in learning strategies across individuals have rarely been addressed, partly owing to the limitations of data gathering and analysis.

Studying differences among individuals requires training and collecting data from multiple animals in a standardized and high-throughput fashion. The training procedures are often time-consuming, requiring several days to many weeks^8,9^, depending on the task. Although there have been advances in training automation, existing systems either require an experimenter to move animals from the home cage to the training apparatus^14–16^, or training animals within their own cages^17–19^. The former introduces additional sources of variability^20,21^, and the latter precludes tasks that require a large training arena. Following data acquisition, the analysis of behavior aims at understanding the learning process. Present approaches tend to focus on the averaged performance over many trials^22^. However, changes in behavior may happen at a single trial, and thus the modeling of behavior should similarly offer a time resolution of single trials to assess each animal’s individual approach to learning.

To address these challenges, we present Mouse Academy, an integrated platform for automated training of group-housed mice and analysis of behavioral changes in learning a decision-making task. We designed hardware that makes use of implanted radio frequency identification (RFID) chips to identify each mouse, and guides the animal into a behavior training box. Synchronized video recordings and decision-making sequences are acquired during animal learning. To analyze the decision-making sequences, we developed an iterative generalized linear model (GLM). This model makes a prediction of the animal’s choice in each trial and gets updated based on the animal’s actual choice. This iterative GLM model achieves a prediction accuracy of ~80%, and also reveals the decision-making strategy of the animal and how it changes over time. To analyze the animal’s behavior during the task in greater detail, we computed the movement trajectories of each mouse. These trajectories allowed us to perform an unsupervised analysis of each animal’s behavior, and discover individual traits of behavioral learning that were not apparent from the simple choice sequences.

## RESULTS

The Mouse Academy platform consists of three components (**Fig. 1**): an automated RFID sorting and animal training system, an iterative GLM to analyze decision-making sequences, and behavior assessment software that analyzes animal trajectories computed from video data.

**Fig. 1:**
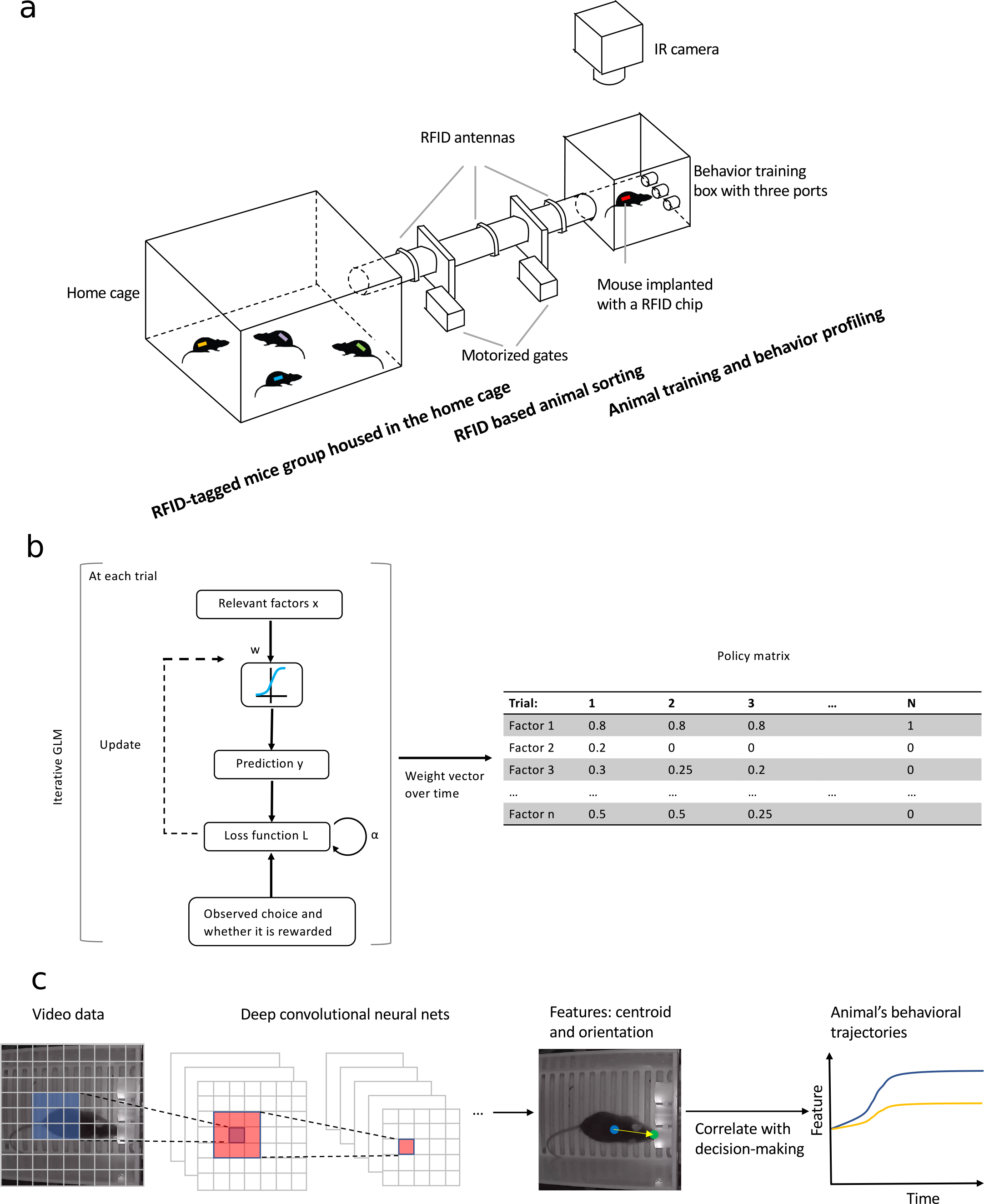
Components of Mouse Academy. **(a)** An automated RFID sorting and animal training system. Mice implanted with RFID chips are group-housed in the home cage. The RFID sorting system identifies each mouse by its implanted chip. One animal at a time gains access to a behavioral training box. As the animal learns a task, its decision sequences and video recordings are acquired. **(b)** An iterative generalized linear model. For each trial, the model predicts the animal’s choice based on the relevant factors and then evaluates the difference from the actual choice. This difference, after temporal weighting, is fed back to the loss function, which gets minimized by updating the weights of the input factors. The model produces a policy matrix in which the rows indicate the weights of the relevant factors and the columns are the trials. **(c)** An automated behavior assessment program using deep convolutional neural networks to extract the location and pose information of an animal.

### Automated RFID sorting supports individual training programs

We designed the equipment in the following manner (**Fig. 1a**): RFID-tagged mice are grouped in a common home cage where food and bedding is supplied. The home cage connects to a behavior training box through a gated tunnel. The gates are controlled by a home-made RFID animal sorting system^23^: three RFID antennas are placed along the tunnel, with one near the home cage, one near the training box and one between the two; the motorized gates are placed between the RFID sensors, separating the tunnel into three compartments. An Arduino microcontroller integrates information from the RFID readers to open and shut the gates, allowing only one animal at a time to pass through the tunnel (**Supplementary Figs. 1a, 1c, 1d** and **Supplementary Videos 1-4**). The behavior box is outfitted with three ports, each of which contains a photo-transistor to detect snout entry, a solenoid valve to deliver water reward, and a light emitting diode (LED) to present visual cues. To maintain a controlled environment, the training box is isolated from the outside by a light- and sound-proof chamber (**Supplementary Fig. 1b**).

Once a mouse enters the training box, a protocol is set up to train the mouse to perform a certain task. In the experiments reported here, the animal must nose-poke the center port to initialize a trial and then hold the position for a short period. Visual or auditory stimuli are delivered, and based on these stimuli, the animal must choose to poke one of the side ports. If the correct response is chosen, the animal gets water reward from a lick tube in the response port, otherwise a timeout punishment is applied. This training process is controlled by Bpod, an Arduino microcontroller that interfaces with the three ports. Data from the response ports as well as video recordings from an overhead camera are acquired simultaneously as the animal is trained.

The entire apparatus is orchestrated by a master program that coordinates the RFID sorting device, the Bpod system, synchronized video recording, data management and logging (**Supplementary Fig. 1e**). The program monitors the amount of water each animal consumes per day and regulates the time each animal can spend in the training box per session. In addition, the software updates the training protocol for each animal based on its performance, for example switching to a harder task once a simpler one has been mastered (**Supplementary Fig. 2**). This lets each animal learn at its own pace.

The apparatus can be assembled at a materials cost of $1500-2500, with the cheaper option using a Raspberry Pi computer as the controller (**Supplementary Fig. 1f** and **Supplementary Table 1**). Compared with designs in which each animal is automatically trained in its own home cage^15,17^, the system saves considerable space. Because housing and training are independent modules, the same system can be used for diverse training environments.

We tested the automated RFID sorting and animal training system by training group-house mice to learn a variety of decision-making tasks, following similar procedures as reported previously^11,12^ (**Supplementary Fig. 2** and Online methods). The training period lasted 28 days, with up to five mice in the common home cage. Each animal occupied the training box for 3-4 hours per day (13-15% of the 24 hours) throughout the entire training period (**Figs. 2a**, **2b** and **Supplementary Fig. 3**). For a sample cohort of four animals trained in sessions of 90 trials each, we found that the behavior box was occupied most of the time, with brief empty intervals of <10 min (**Figs. 2c, 2d and 2e**). Each animal was trained for over 900 trials (10 sessions), and consumed more than 1.9 mL of water per day (**Fig. 2f**). Interestingly there was no circadian pattern to the animals’ training activity, even though the setup was illuminated on a daily light cycle (12 h on / 12 h off) (**Fig. 2g**). As observed previously, it appears that animals working for a goal can avoid circadian modulation of the locomotor pattern^24,25^.

**Fig. 2:**
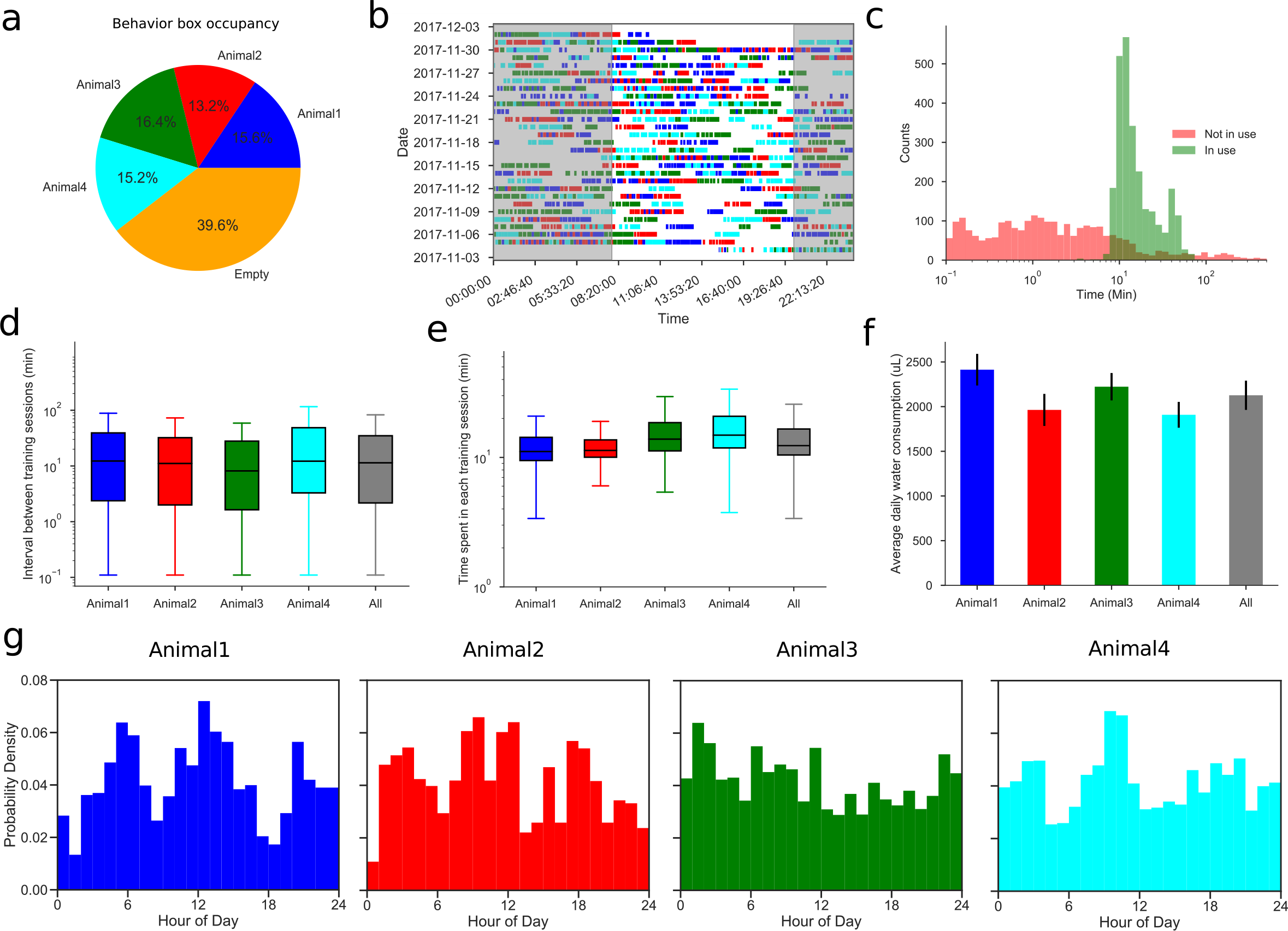
Performance of the automated training system on a sample cohort. **(a)** Fraction of time the behavior box was occupied by each of the four animals. **(b)** Activity trace of each animal in the behavior box for the entire training period of 28 days. Shadow indicates the dark cycle from 8pm to 8am. **(c)** Distribution of time intervals during which the behavior box is occupied or empty. **(d)** Box plot of intervals between each animal’s sessions (median, quartiles, and range). **(e)** Box plot of the time spent in a session for each animal. **(f)** Averaged daily water consumption of each animal. Error bars indicate standard errors. **(g)** Circadian histograms of each animal’s activity in the behavior box.

### A generalized linear model accurately predicts decision-making during training

In a decision-making task, an animal is asked to associate distinct stimuli with distinct responses. Although this is the ultimate goal, during learning, it is often observed that the animal begins by basing its decisions on unrelated input variables and gradually switches to using the stimulus variables that actually predict reward. We define a policy as a mapping of these variables to the animal’s decisions. A fundamental goal in the study of learning is to infer what policy the animal follows at any given time and to determine how the policy evolves with experience.

We applied a generalized linear model (GLM) to map factors relevant to the animal’s decision-making to its choices through logistic regression. A common way to build such a GLM is by fitting data of an entire session^16,26^. However, this loses resolution in single trials within the session. During learning, a change of policy can happen at each trial. Thus, we developed the model to make trial-by-trial choice predictions based on various factors the animal might plausibly use. The model works in an iterative two-step process (**Fig. 1b**). In the prediction step, the model makes a prediction for the next decision based on the input factors. Once the outcome of the animal’s decision is observed, an error term between the model’s prediction and the observation is computed. This error, after weighting by a reward factor and a temporal discount factor, is fed back to the loss function. In the update step, the model is updated by minimizing the regularized loss function. This iteration happens after every trial. The temporal discount factor accounts for the possibility that the most recent trials impact the current decision more than remote trials. The reward factor accounts for the fact that water rewards and timeout punishments may have effects of different magnitude on the updates of the animal’s policy.

We illustrate the utility of this model by fitting results from an easy visual task, in which one of the two choice ports lights up to indicate the location of the reward, and the optimal policy is to simply poke the port with the light (**Supplementary Fig. 2a, 2a’ and 2a’’**). All the mice eventually reached a >83% performance level, comparable to what mice achieve in similar tasks^19,27^. The GLM makes a prediction for the outcome of each trial based on a weighted combination of several input variables: the current visual stimulus, a constant bias term, and three terms representing the history of previous trials (**Fig. 3a**). These inputs from a previous trial include the port choice, whether that choice was rewarded, and a term indicating the multiplicative interaction between the choice and reward (Choice × Reward). This term supports a strategy called win-stay-lose-switch (WSLS), which chooses the same port if it was rewarded previously and the opposite one if not. Since a GLM cannot multiply two inputs, we provided this interaction term explicitly. Each of the above terms has a weight coefficient that can be positive or negative. For instance, a positive weight for the visual stimulus supports turning towards the light, and a negative weight away from the light.

**Fig. 3:**
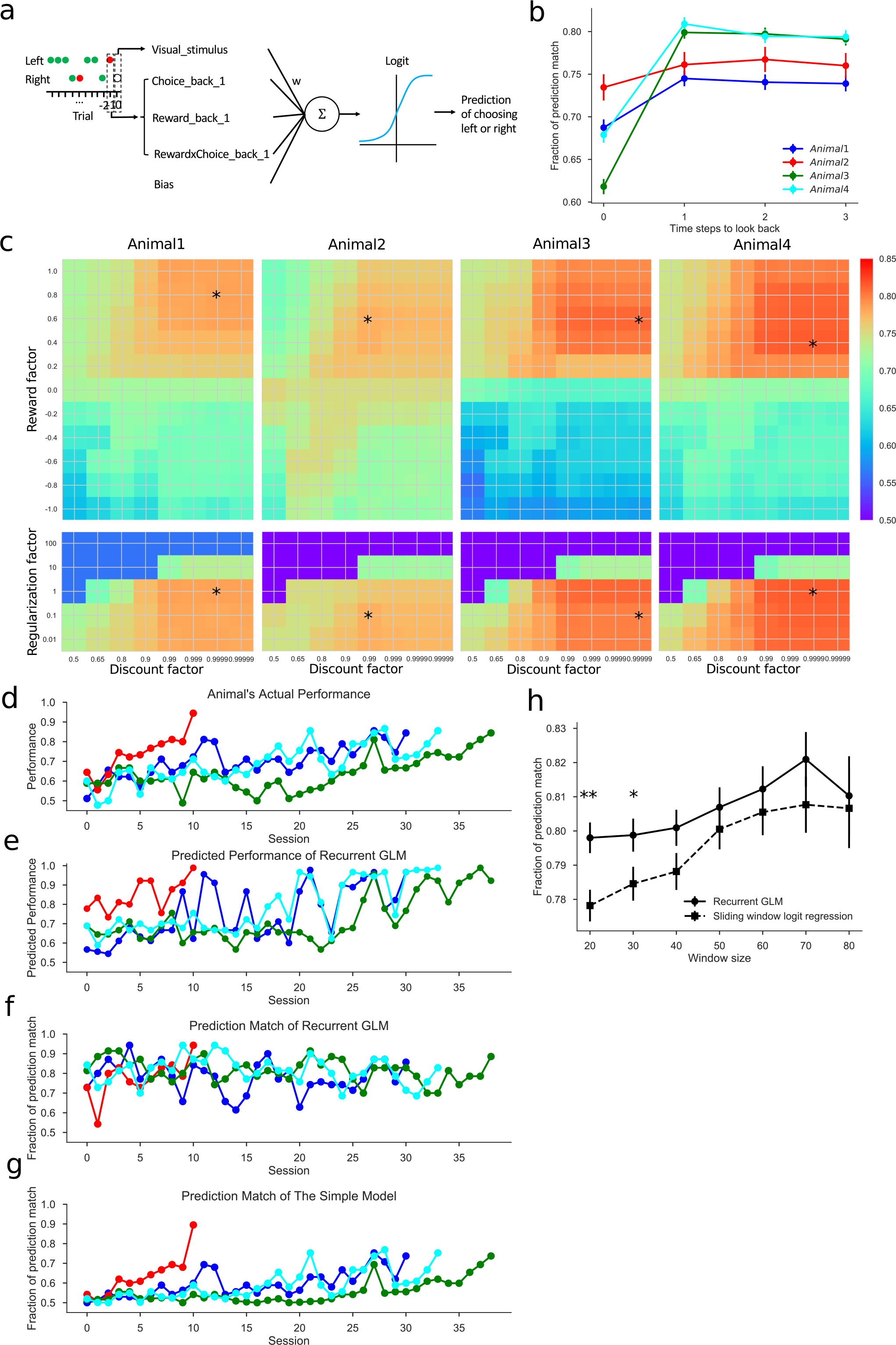
Iterative generalized linear model and its prediction accuracy. **(a)** Illustration of the GLM as applied to a visual discrimination task. The model’s prediction is based on the output of a logistic function whose input is the weighted sum of a visual stimulus term, a bias term, and three history-dependent terms. The stimulus can be on the left or right and the choice can be rewarded (consistent with the stimulus, indicated by a green dot) or unrewarded (opposite to the stimulus, indicated by a red dot). **(b)** Selection of the history-dependent terms based on the model prediction accuracy. Error bars indicate standard errors. **(c)** Hyperparameters for each of the animals: reward factor, discount factor, and regularization factor. The optimal values are marked with a star. **(d)** The actual performance of each animal over time in the visual task. **(e)** Performance as predicted by the GLM. **(f)** Fraction of choices predicted correctly by the GLM. **(g)** Fraction of choices predicted correctly by a simple model based on the animal’s average performance in the task. **(h)** Fraction of predictions matched by the iterative GLM and the sliding window logistic regression model. Error bars indicate standard errors. **, * indicate P < 0.01, 0.05. Random prediction would give 50% match.

To determine the extent of trial history that affects the animal’s behavior, we fitted the model to the response data including history-dependent terms up to three previous trials. We found that only the immediately preceding trial had an appreciable effect on the prediction accuracy, and thus restricted further analysis to those inputs (**Fig. 3b**). The model also has three hyperparameters (the temporal discount factor *α*, the reward factor *r*, and the regularization factor *λ*), and we optimized them for each animal by grid search. We found that each animal had a different set of hyperparameters, reflecting differences in the learning process across individuals (**Fig. 3c**). Among the four sample mice, Animal 2 had the lowest temporal discount factor, suggesting that it weighed recent trials more heavily and updated the policy more quickly. Indeed, this is the animal that learned the fastest among the four (**Fig. 3d**).

Predictions from the iterative GLM matched ~80% of the animals’ actual choices (**Fig. 3f**), and the predicted accuracy of each animal captured the actual fluctuations of its learning curve (**Fig. 3e** and **Supplementary Fig. 4**). We compared the performance of the GLM with two other modeling approaches (Online methods). The first model was fit to the animal’s average performance in the task; its trial-by-trial match of the animal’s actual choices was only ~59% (**Fig. 3g**). The second model was a logistic regression fitted to data in a sliding window of *N* trials. This sliding window model performed worse than the iterative GLM when the window size was small (*N* = 20 and 30 trials, **Fig. 3h**); for larger windows the performance was comparable. Overall, the iterative model is advantageous because it makes predictions online as every trial occurs and adapts dynamically to the growing data set.

### Individual learning policies can be inferred from iterative GLM fitting

The iterative GLM serves to infer what policy the animal follows in making decisions. The linear weight of each input term reflects its relative importance for the decision. By following this weight vector across trials one obtains a policy matrix that documents how the animal’s policy changes during learning (**Figs. 1b** and **4c**). To test that the model can correctly capture a time-varying policy, we simulated decision-making data from a ground truth policy that changed at a certain frequency, including a certain level of noise in the behavioral output (**Fig. 4a**). Over a wide range of policy change frequencies and noise levels, the GLM was able to capture the ground truth policy (**Figs. 4a and 4b**). In addition, different values of policy change frequency and noise levels led to different sets of hyperparameters fitted from the model, showing that the GLM can adapt to individuals with diverse learning characteristics (**Supplementary Figs. 5a-e**).

**Fig. 4:**
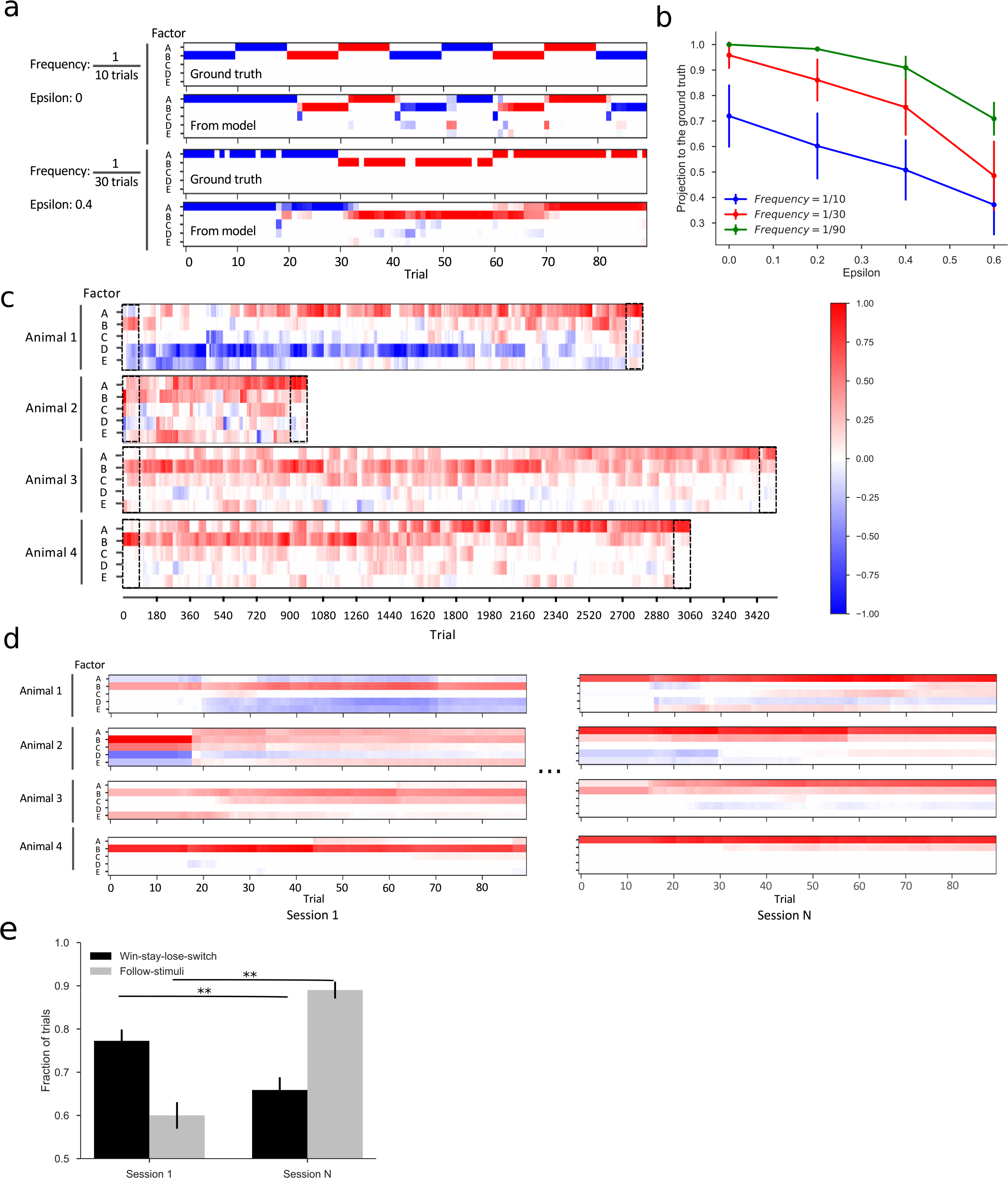
Interpretation of policies during learning. **(a)** Policy vectors recovered by the iterative GLM capture the ground truth policies. The policy matrix plots in each trial (horizontal) the weights associated with each of 5 factors (vertical), encoded with a color scale (see panel c). The factors are: A = Visual_stimulus, B = Choice × Reward_back_1, C = Choice_back_1, D = Reward_back_1, E = Bias. Two examples are shown of ground truth policies used to simulate data and the corresponding trial-by-trial estimates from the GLM. Blanks in the ground truth matrix indicate instances where the simulated choice is opposite to the policy. **(b)** Similarity between the recovered policy and the ground truth, measured by the cosine between the two policy vectors. Error bars indicate standard deviation. **(c)** Policy matrices recovered for the four animals show distinct individual learning processes. Dashed rectangles highlight the first and last sessions of each animal, as enlarged in **d**. **(d)** Recovered policy matrices for the first and last sessions of each animal. **(e)** Fraction of trials explained by two candidate policies (win-stay-lose-switch and following the stimuli) in the first and last sessions. Error bars indicate standard errors. ** indicates P < 0.01.

We then recovered the policy matrix of each animal from the GLM fits. All four animals started with the non-optimal policy of WSLS. Subsequently each animal followed its own learning process (**Fig. 4c**): Animal 2 had a clear bias towards the right port at the beginning but it rapidly found the optimal policy of following the light. The other three animals were slower learners. Animal 3 and Animal 4 followed similar processes to converge to the optimal policy. Animal 1 was distinct from the others. At the early stages, it had a strong bias towards the left port and it made decisions based on whether the previous choice was rewarded.

We further validated the transition between policies during learning by analyzing the first and last sessions of each animal and counting how many choices could be explained by each policy (**Fig. 4d**). Indeed, we found a clear switch from the (non-optimal) WSLS policy to the (optimal) stimulus-based policy (**Fig. 4e** and **Supplementary Fig. 5f**). The animals might have been biased towards the WSLS strategy by a shaping method we used during training, which offered the animal a repeat of the same stimulus every time it made a mistake (Online methods). To test whether these correlations in the trial sequence influenced the final policy we performed two additional analyses. First, we only included trials following a correct trial, and performed logistic regression on these trials for each session. This analysis showed that at least on these trials, all the animals based their decisions on the light stimulus by the end of learning (**Supplementary Fig. 6a**). Second, we compared the error rate on trials following an incorrect choice with that following a correct one. We found no significant difference between the two error rates during the last session (**Supplementary Figs. 6b and 6c**), suggesting that the animals treated these two types of trials identically.

### Automated movement tracking reveals fine structure of behavioral responses

Thus far the report has focused on the animal’s responses only as sensed by the nose pokes into response ports. The GLM fits of those responses already revealed differences in policy across individuals. To gain further insight into these individual preferences, it is essential to track each animal’s behavior along the way from stimuli to responses^10^. We thus employed computer vision software to automatically, quantitatively and accurately assess each animal’s behavior during decision-making. Having compared several tracking algorithms, we eventually used DeepLabCut^45^, a deep-learning based program, because it is easy to use and accurate in identifying body landmarks (**Figs. 1c** and **5a**). We identified two body landmarks: the nose and the centroid of the animal, and further calculated the orientation as the angle of the line connecting the centroid and the nose (**Fig. 5a, Supplementary Video 5**).

**Fig. 5:**
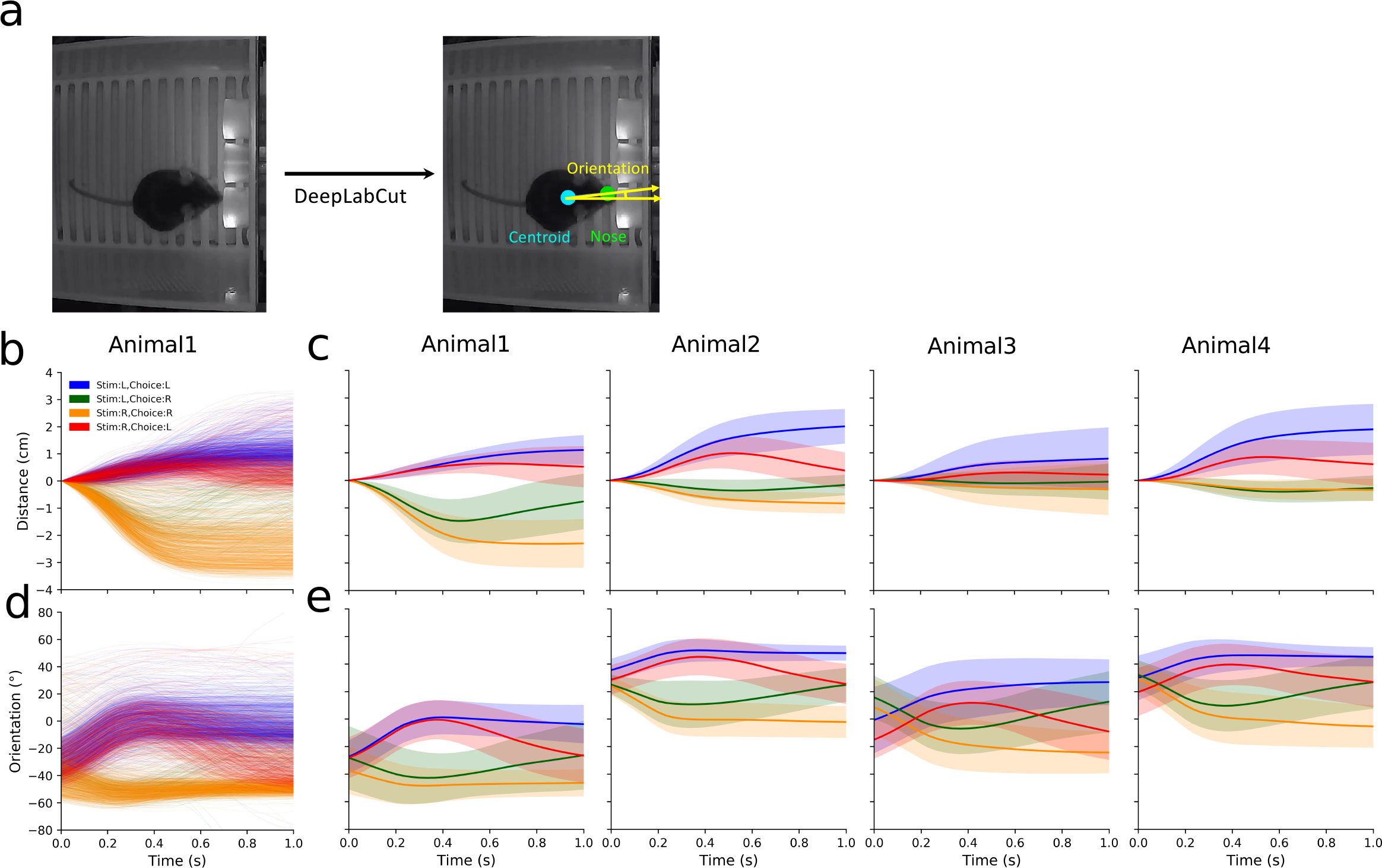
Supervised analysis using features extracted by automated behavior assessment. **(a)** DeepLabCut extracts the centroid and the orientation as the angle between the horizontal axis and the line connecting the centroid and the nose. **(b)** Centroid distance along the left-right axis vs time during the movement, for animal 1. The starting position is set to zero, positive values indicate movement to the left, negative to the right. The 4 trial types are indicated by different colors. **(c)** Average centroid trajectory for each animal. Shaded region indicates standard error. **(d-e)** Orientation vs time, displayed as in panels b-c. Positive angle points to the left, negative to the right.

To illustrate use of these behavioral trajectories, we focus on the period of the visual choice task where the animal reports its decision: from the time it leaves the center port to when it pokes one of the side ports. The trials fall into four groups based on location of the stimulus and the response. As expected, the trajectories of position and orientation clearly distinguish left from right choices (**Figs. 5b and 5d**). Interestingly, the trajectories also reveal whether the decision was correct: On incorrect decision, the trajectories reversed direction after ~0.5 s, because the animal quickly turned back to the center after finding no reward in the chosen port (**Figs. 5c, 5e** and **Supplementary Video 5**). A linear kernel support vector machine (SVM), trained to predict the category of each trial from a 1 s trajectory, was able to correctly distinguish correct and incorrect choices with an accuracy of over 90% (**Supplementary Fig. 7**). In addition, many of the trajectories were highly asymmetric and again revealed differences across individuals. For instance, Animal 2 and Animal 4 started from a location close to the right port, Animal 1 closer to the left port (**Fig. 5c**). This asymmetry correlates with the bias revealed by the iterative GLM: each animal prefers to select the port closer to its body location.

### Unsupervised behavioral analysis reveals moments of hesitation

Whereas the supervised learning discussed above relies on prior classification of stimuli and responses, an unsupervised analysis has the potential to discover unexpected structures in the animal’s behavior^28^. We thus performed an unsupervised classification of the behavioral trajectories.

After subjecting all the trajectories of a given animal to principal component analysis (PCA) we projected the data onto the top three components, which explained over 95% of the variance (**Figs. 6a and 6b**). Importantly, without any labels from trial types, these three PCs captured meaningful features that differentiated the animal’s responses. The first PC separated movements to the left from those to the right. The third PC captured the turning-back behavior after an incorrect choice. The second PC captured different baseline positions. Each animal has its own preference for a baseline position somewhere off the midline of the chamber (**Supplementary Fig. 8a**).

**Fig. 6:**
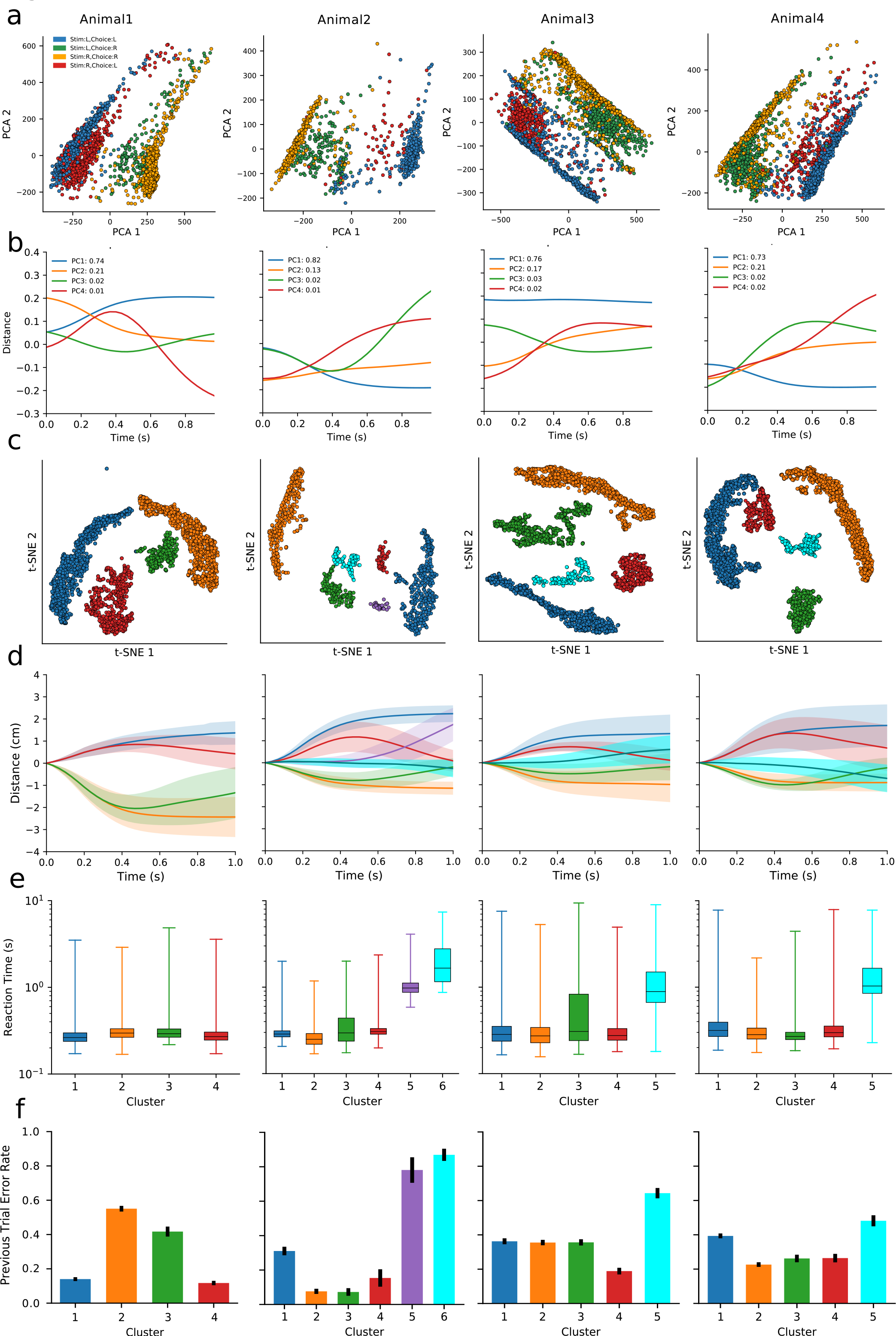
Unsupervised analysis of the behavior trajectories. **(a)** Principal component projections onto PC1 and PC2 of the centroid-vs-time trajectories from Fig 5. The 4 trial types are indicated by different colors. **(b)** The centroid trajectories corresponding to the first four principal components (PCs). The variance explained by each PC is shown in the plot legend. **(c)** Clustering trials by their trajectories using t-SNE analysis. Distinct clusters are marked with different colors for use in subsequent panels. **(d)** Averaged centroid distance vs time for each cluster, plotted as in Fig 5b. **(e)** Box plot of the reaction time for each cluster. **(f)** The error rate on the preceding trial for each cluster. Error bars indicate standard errors.

We also projected the trajectories into 2 dimensions using a non-linear embedding method, t-distributed stochastic neighbor embedding^29,30^ (t-SNE). Unlike PCA, this graph prioritizes the preservation of local structures within the data instead of the global structure^30^. In the t-SNE space the trajectories formed clear clusters (**Fig. 6c**). Most of the clusters are dominated by one of the decision categories (**Fig. 6c** and **Supplementary Fig. 8b**). Interestingly, we found clusters in Animals 2, 3, and 4, in which the centroid trajectories were flat, unlike the trajectories of the four decision categories (**Fig. 6d**), suggesting that animals hesitated in these trials and made decisions only after a delay. Indeed, in trials flagged by these clusters, the animals had longer reaction times (**Fig. 6e**). Furthermore, such hesitating responses were more common following an incorrect trial (**Fig. 6f**); they may reflect a behavioral adjustment to prevent further mistakes^31^.

## DISCUSSION

Despite the fact that rodents can be trained to perform interesting decision-making tasks^7–10^, the learning progress of individual animals has rarely been addressed. Doing so requires training and observing many animals in parallel under identical conditions, and the ability to analyze the decision policy of each animal on a trial-by-trial basis. To meet these demands, we developed Mouse Academy, an integrated platform for automated training and behavior analysis of individual animals.

We demonstrate here that Mouse Academy can train group-housed mice in an automated and highly efficient manner while simultaneously acquiring decision-making sequences and video recordings. Automated animal training has been of great interest in recent years and efforts have focused on two directions. In one design, multiple animals are trained in parallel within stacks of training boxes. This requires a technician to transfer animals from their home cages to the behavior boxes^14–16^. Such animal handling has been reported to introduce additional variability^20,21^, and even the mere presence of an experimenter can influence behavioral outcomes^32^. Thus, eliminating the requirement for human intervention, as in Mouse Academy, likely reduces experimental variation. In another design, a training setup is incorporated within the animals’ home cage^17–19^. By contrast, Mouse Academy separates the functions of housing and training, and that modular design allows easy adaptation to a different purpose. For instance, one can replace the 3-port discrimination box with a maze to study spatial navigation learning^33,34^, or with an apparatus for training under voluntary head-fixation^35^. In each case, a single training apparatus can serve many mice, potentially from multiple home cages.

To understand how an animal’s decision-making policies change in the course of learning, we developed a trial-by-trial iterative GLM. The evolution of the model is similar to online machine learning^36^ in which the data are streamed in sequentially, rather than in batch mode. The linear nature of the model supports a straightforward definition of the animal’s decision policy, namely as the vector of weights associated with different input variables. In addition, the simple linear structure allows rapid execution of the algorithm, which favors its use in real-time closed-loop behavior experiments. The model also allows several parametric adjustments. One specifies how much the recent trials are weighted over more distant ones in shaping the animal’s policy. Another rates the relative influence of reward versus punishments. Fitting these parameters to each animal already revealed differences in learning style. This model can have a broader use beyond mouse decision-making, for instance to track the progress of human learners from their answers to a series of quizzes^37^.

Finally we analyzed behavioral trajectories of individual animals. The behavioral trajectories can reveal intricate aspects of the animal’s decision process that are hidden from a mere record of the binary choices. The large data volume again calls for automated analysis, and both supervised machine learning methods^28,38,39^ and unsupervised classification^28–30,40^ have been employed for this purpose. Unsupervised analysis is not constrained by class labels, and can identify hidden structure in the data in an unbiased manner. In the present case, we discovered a motif wherein the animal hesitates on certain trials before taking action.

Mouse Academy can be combined with chronic wireless recording^41,42^, to allow synchronized data acquisition of neural responses. Researchers can seek correlations between neural activity and the policy matrix or even the behavioral trajectories. This will open the door to a mechanistic understanding of how neural representations and dynamics change in the course of animal learning.

## ONLINE METHODS

### Animals

Subjects were C57BL/6J male mice aged 8-12 weeks. All experiments were conducted in accordance with protocols approved by the Institutional Animal Care and Use Committee of the California Institute of Technology.

### Hardware setup

The hardware setup comprises a behavioral training box, an engineered home cage, and a radio frequency identification (RFID) sorting system, which allows animals to move between the home cage and the training box. These components are coordinated by customized software.

The design file for the behavior box was modified from that of Sanworks LLC (https://github.com/sanworks/Bpod-CAD) using Solid-Works computer-aided design software and the customized behavioral training box was manufactured in the lab. The behavior box is controlled by a Bpod state machine (r0.8, Sanworks LLC). To monitor the animal’s behavior, an IR webcam (Ailipu Technology or OpenMV Camera M7) is installed above the behavior box. The behavior box and the webcam are placed within a light- and sound-proof chamber. The chamber is made of particle board with walls covered by acoustic foam. A tunnel made of red plastic tubes connects the behavior box to a home cage (**Supplementary Fig. 1b)**.

For the RFID access control system, an Arduino Mega 2560 microcontroller is connected with three RFID readers (ID-12LA, Sparkfun) with custom antenna coils spaced along the access tunnel. The microcontroller controls two generic servo motors fitted with plastic gates to grant individual access to the training box (**Supplementary Fig. 1a)**.

The microcontroller identifies each animal by its implanted RFID chip and permits only one animal to go through the tunnel connecting the home cage and the behavioral training box (**Supplementary Fig. 1c)**. It also communicates the animal’s identity to a master program running on a PC or Raspberry Pi (Matlab or Python). The master program coordinates the following programs: Bpod (https://github.com/sanworks/Bpod), synchronized video recording, data management and logging. A repository containing the design files, the firmware code for the microcontroller, and the software can be found in https://github.com/muqiao0626/Mouse_Academy.

### Behavior training

The training procedures of mice to perform a selective attention task are similar to those previously reported^11,12^. Mice were water restricted for seven days before training, and habituated in the automated training system to collect reward freely for several sessions. Then the mice were trained in sessions, each of which was made of 90 trials, to collect water rewards by performing two alternative forced choice tasks. Briefly, the animal had to nose-poke one of two choice ports based on the presented stimuli. If the decision was correct, 10% sucrose-sweetened water (3 μL) was delivered to the animal. For incorrect responses, the animal was punished with a five-second timeout. Following an incorrect response, the animal was presented with the identical trial again; this simple shaping procedure helps counter-act biases in the behavior.

Over 28 days of training the animals learned increasingly complex tasks, from visual discrimination to a two-modality cued attention switching task^11,12^. The training progressed through six stages (**Supplementary Fig. 2**):

1. A simple visual task: In this task, the animal initiates a trial by poking the center port and holding the position for 100 ms. Then either the left or right side port light up briefly until the animal moves away from the center port. The animal must then poke one of the two side ports within the decision period of 10 s. Choice of the port flagged by the light leads to a water reward, and choice of the other port leads to a time-out period during which no trials can be initiated. Data presented in the main text are from this stage of training only.
2. A simple auditory task: As Stage 1, except that the stimulus was white noise sound either the left or the right side to flag the reward port.
3. A cued single-modality (visual or auditory) switching task: Blocks of 15 trials consisting of single-modality (visual or auditory) stimulus presentation. Each block was like stages 1 or 2, except that the trial type was indicated by a 7 kHz (visual) or 18 kHz (auditory) pure tone.
4. A cued single- and double-modality switching task: Like stage 3, but distracting trials were introduced in which both visual and auditory stimuli were present, but only one of the modalities was relevant to the decision. The relevant modality was again indicated by the pure tone cues. In repeating blocks, four types of trials were presented: a. five visual-only trials; b. ten ‘attend to vision’ trials with auditory distractors; c. five auditory-only trials; d. ten ‘attend to audition’ trials with visual distractors. During the training, the time that the animal had to hold in the center port was gradually increased to 0.5 s, and the duration of the stimuli was gradually shortened to 0.2 s.
5. A cued double-modality switching task: Like stage 4 except that the single-modality trials were removed, and the block length was gradually shortened to three trials.
6. A selective attention task: Like stage 5, but the block structure was abandoned and all eight possible trial types were randomized: (audition vs vision) × (sound left or right) × (light left or right).

### Iterative generalized linear model

We modeled the animal’s choice probability by a logistic regression. At each trial number *t*, the choice probability is defined as

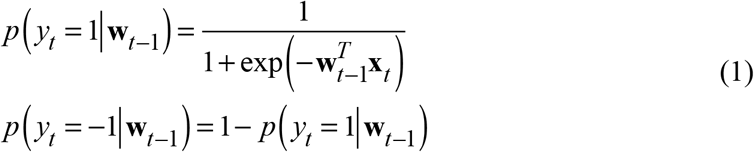

where *y_t_* indicates the binary choice of the animal (1 = right, –1 = left), **x**_*t*_ is the vector of input factors on trial *t*, and **W**_*t*-1_ is the vector of weights for these factors obtained from fitting up to the preceding trial.

The prediction 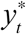 for the animal’s choice is simply that with the higher model probability:

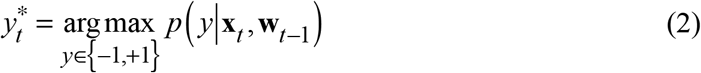

After observing the animal’s actual choice *z*_*t*_, the cross-entropy error *E*_*t*_ between the observation and model prediction is calculated as

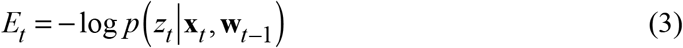

We weight the error term by a reward factor *R*_*t*_, and apply exponential temporal smoothing to get the loss function *L*_*t*_:

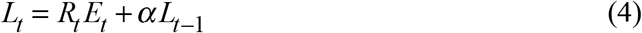

where *α* is the smoothing discount factor accounting for the effect that distant trials have less impact on decision-making than immediately preceding trials, and *R*_*t*_ is defined as

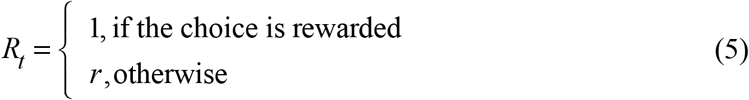

The values of *R*_*t*_ for rewarded and unrewarded trials may be different, accounting for the fact that rewards and punishments may have different effects on learning. For each time point, the weights in the model are determined by minimizing the loss function subject to L1 (lasso) regularization, namely

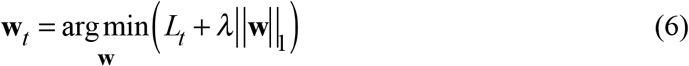

Then **w**_*t*_ is used for prediction of the next trial. For subsequent analysis, we only used predictions starting at the 15th trial. The three hyperparameters for the temporal discount factor *α*, the reward factor *r*, and the regularization factor *λ* were selected by grid search.

To fit the decision-making sequences of the simple visual task, we included the following terms in the input vector **x**_*t*_ :

1. Visual_stimulus: +1 = light on right, –1 = light on left.
2. Bias: A constant value of +1. The associated weight determines whether the animal favors the left (negative) or the right (positive) port.
3. Choice_back_n: The choice the animal made *n* trials ago (+1 = right, –1 = left).
4. Reward_back_n: The reward the animal received *n* trials ago (+1 = reward, –1 = punishment).
5. Choice × Reward_back_n: The product of terms 4 and 5. This term corresponds to the win-stay-lose-switch (WSLS) strategy of repeating the last choice if it was rewarded and switching if it was punished.

To determine the extent of history-dependence of the animal’s decisions, we fitted the model including terms 3-5 from up to three previous trials (*n* = 1, 2, 3), and found that only the immediately preceding trial had an appreciable effect on the model’s prediction accuracy. For the subsequent analysis, we therefore included terms 3-5 for the preceding trial (*n* = 1).

We compared the iterative generalized linear model (GLM) with two other models. The first only captures the animal’s average performance over all trials. If the fraction of the correct responses is *z*, then the model simply predicts a correct response with probability *z*, and an error with probability 1–*z*. Thus, the fraction of trials where the prediction matches the observation is *z*^2^ +(1-*z*)^2^.

The second model is a sliding window logistic regression. To make a prediction for trial *t*, we fitted the logistic model presented above (Eqns 1–2) to the preceding *n* trials. The loss function is

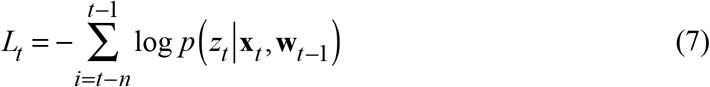

and the weights are again optimized as in Eqn 6.

### Recovering policy matrices from simulated data

To test the model’s ability in recovering policy matrices, we trained the model on data generated from pre-defined ground truth policies. The ground truth policies changed every 10 trials, 30 trials, or 90 trials. Binary choices were simulated with different noise levels using the algorithm ‘epsilon-greedy’: with a probability of epsilon, the simulator made a random choice and with a probability of 1-epsilon it chose the action indicated by the ground truth policy. The noise levels (epsilon values) ranged from 0 to 0.6. The similarity between the recovered policy and the ground truth policy was evaluated by the cosine between the recovered weight vector and the ground truth weight vector.

### Supervised and unsupervised analysis of behavioral trajectories

We annotated two body landmarks, the nose and the centroid, on ~100 frames of the video, and used them to train DeepLabCut^45^. Tested on a separate set of annotated frames, more than 85% the nose positions are inferred within an error radius of 0.25 cm, and more than 85% of the centroid positions are inferred within an error radius of 0.4 cm. From the nose and the centroid, we calculated the orientation as the angle of the line connecting the centroid and the nose. For each trial, the centroid and orientation were extracted for *n* frames (*n* = 30 (1 s) in most cases), thus the data dimension for each trial is 3*n* (the two centroid coordinates and the orientation).

To determine whether the behavioral trajectories contain information about the decision categories, a support vector machine (SVM) with a linear kernel was trained for each decision category. The training set was labelled with the decision category based on information about the visual stimulus and the animal’s choice (for example, “Stim: R, Choice: L” means that the light is on the right and the animal chooses the left port). Performance of the trained SVM was examined by prediction accuracy on the test set, and the F1 score, which is the harmonic mean of precision and recall:

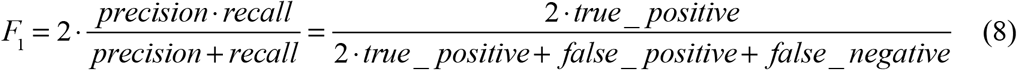

The performance was computed as the average across 10 repeated analyses (**Supplementary Fig. 7**).

We performed a non-linear embedding method, t-distributed stochastic neighbor embedding (t-SNE) analysis as previously described^29,30^. Briefly, the trajectory data of each trial were projected into a 2D t-SNE space. Point clouds on the t-SNE map represented candidate clusters. Density clustering identified these regions. We then plotted trajectories and reaction time distributions to confirm that the clusters were distinct from each other. A repository of the analysis scripts can be found in https://github.com/tonyzhang25/MouseAcademyBehavior.

## AUTHOR CONTRIBUTIONS

M.Q. and M.M. designed the study; M.Q. and S.S. constructed the hardware setup and wrote the controlling software; M.Q. performed experiments and collected data for analysis; M.Q. developed the iterative generalized linear model with input from P.P. and M.M; C.S. implemented animal tracking and provided help in analyzing the mouse trajectories; T.Z. analyzed behavioral trajectories with input from M.Q., P.P. and M.M; M.Q. and M.M. wrote the manuscript with comments from all authors.

## ACKNOWLEDGEMENT

We thank Joshua Sanders for technical assistance in incorporating Bpod into our system. We thank Ann Kennedy for insightful comments and suggestions on analysis of the behavioral trajectories. We thank Oisin Mac Adoha and Yuxin Chen for helpful comments and discussions. This work was supported by a grant from the Simons Foundation (SCGB 543015, M.M. and P.P.) and a postdoctoral fellowship from the Swartz Foundation (M.Q.).

## DATA AVAILABILITY

The datasets analyzed during the current study are available in https://drive.google.com/open?id=1gkPbqGYKPGs7Rx1WNmubQW0dKyYE5YVR

## Supplementary Materials

**Supplementary Table 1.**
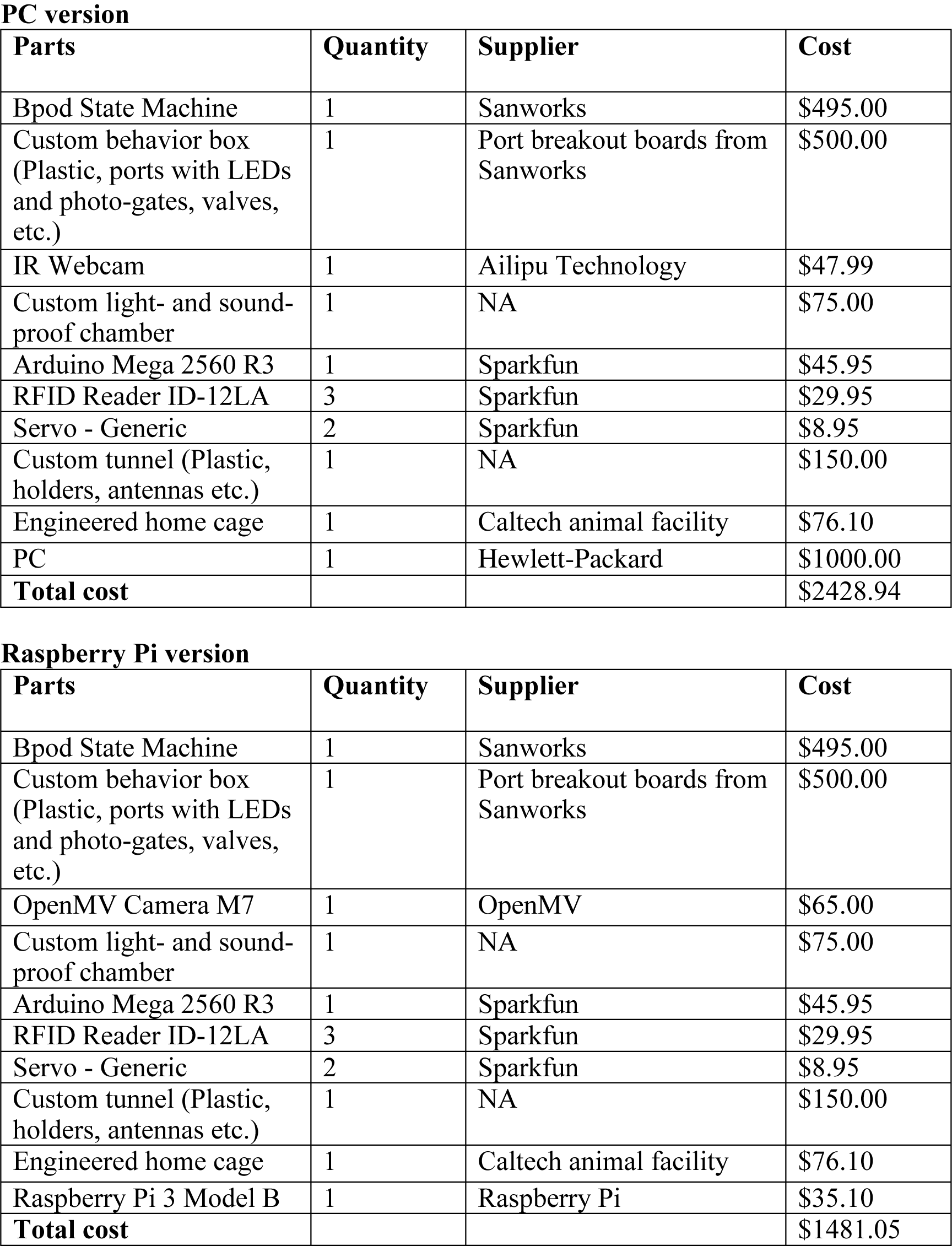
Cost Analysis of Mouse Academy Hardware.

**Supplementary Table 2:** mean averaged precision (mAP) and recall (mAR) at different threshold of Intersection over Union (IoU) (related to mouse detection)

|  | mAP | mAR |
| --- | --- | --- |
| $0.75 < IoU < 0.95$ | 0.81 | 0.82 |
| $IoU = 0.5$ | 0.99 | 0.99 |
| $IoU = 0.75$ | 0.95 | 0.98 |

**Supplementary Fig. 1:**
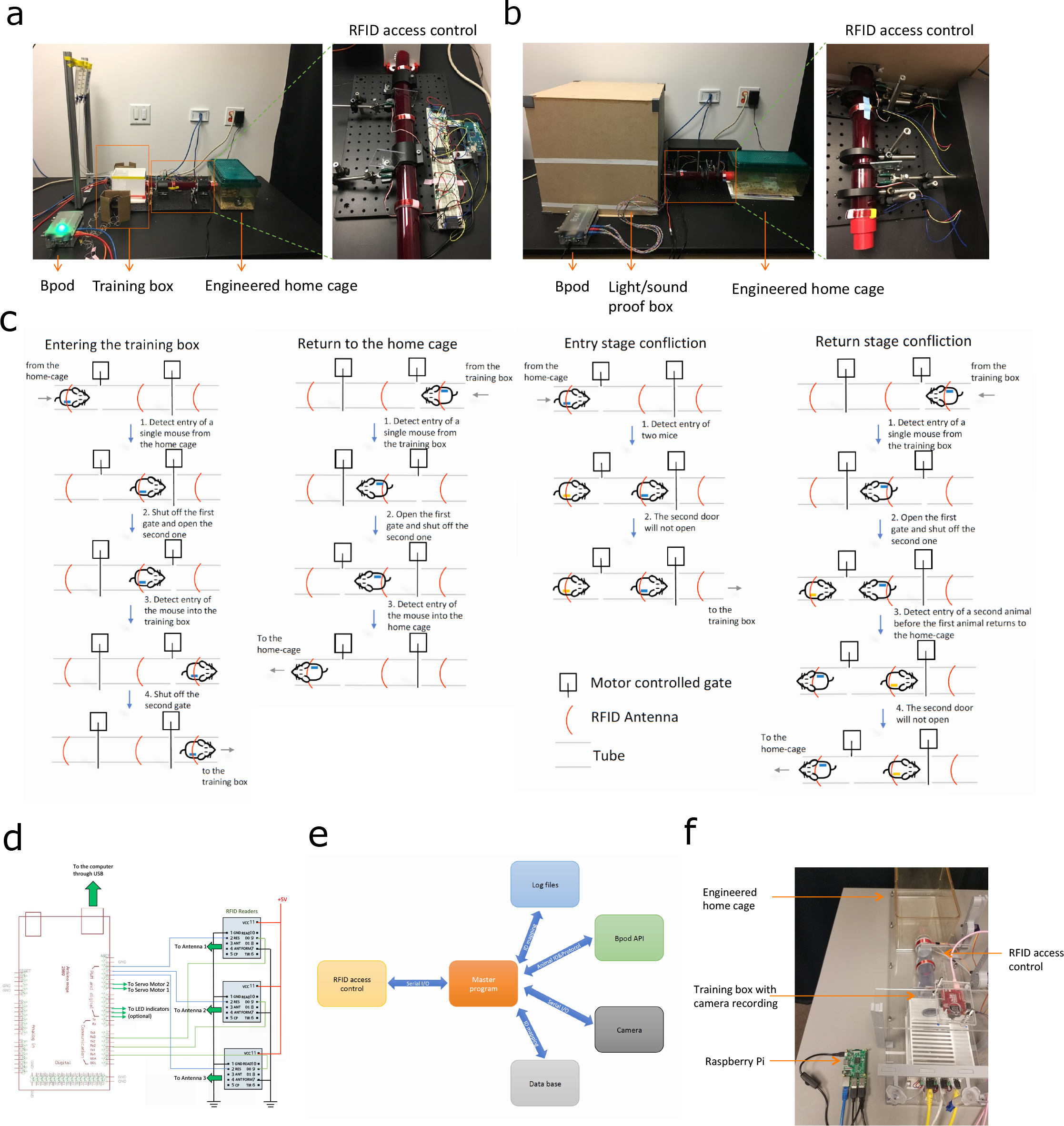
Technical details of the hardware design. **(a-b)** Side view of the setup (**a**) packed into a light- and sound-proof box (**b**). **(c)** RFID sorting process. For an animal to enter the behavior box, only when the left and the middle detectors detect the same RFID chip, the left gate is closed and the right gate is open so that the animal can access the behavior box. For an animal to return to the home cage, only when the right and the middle detectors detect the same RFID chip, the right gate is closed and the left gate is open so that the animal can go back to the home cage. In the entry and the return processes, if the left and the middle detectors detect different RFID chips, the animals have to leave the tube and the detectors get reset afterwards. **(d)** Schematic of RFID access control circuit. **(e)** Schematic of the software controlling the devices. A master program receives input from the RFID sorting device and controls four other modules including Bpod, synchronized video recording, data management, and logging. **(f)** Top view of a Raspberry Pi version of the setup.

**Supplementary Fig. 2:**
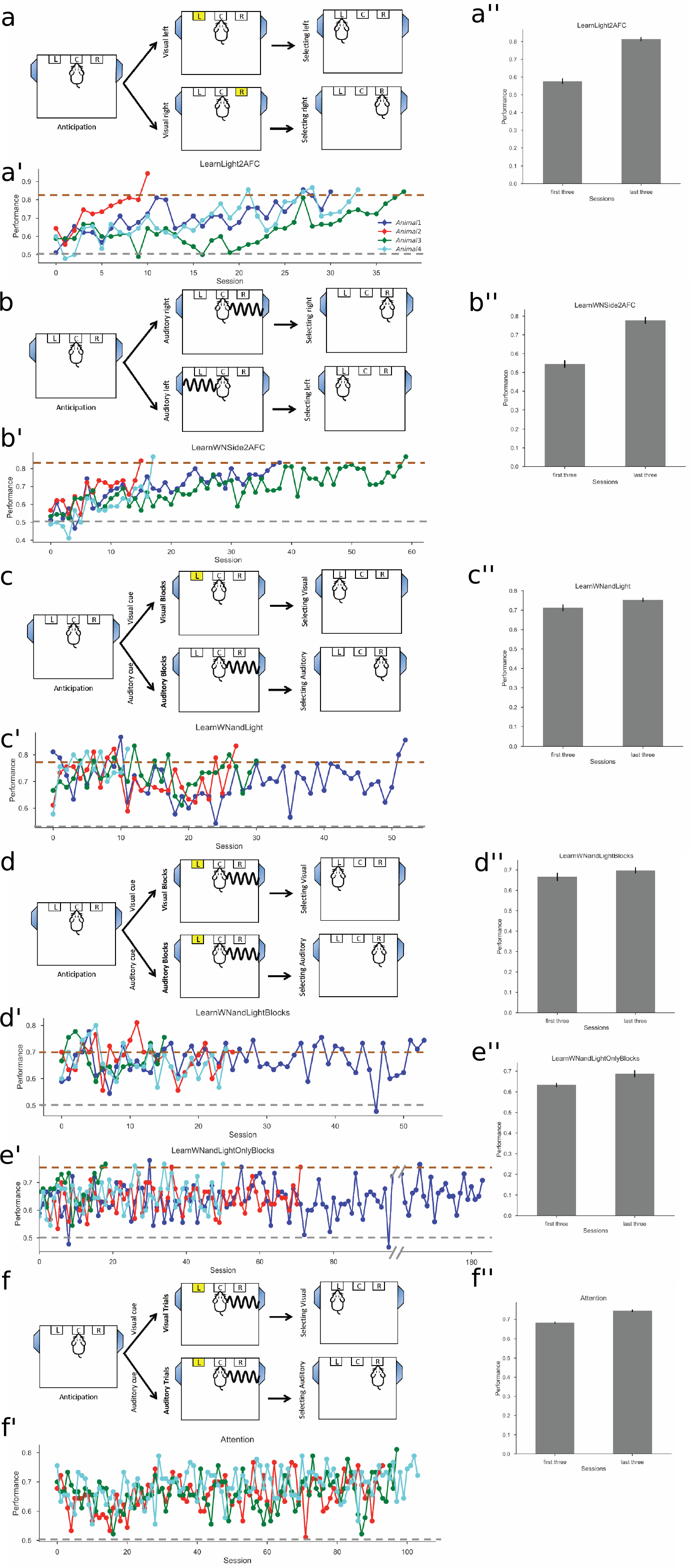
Illustration of training procedures. Training proceeds through six stages (Online methods). The design, learning curves, and animal performance of the simple visual task **(a, a’, a’’)**, the simple auditory task **(b, b’, b’’)**, the cued single-modality (visual or auditory) switching task **(c, c’, c’’)**, the cued single-(visual or auditory) and double-modality (attend to vision or audition) switching task **(d, d’, d’’)**, the cued double-modality (attend to vision or audition) switching task **(e’, e’’)**, and the final selective attention task **(f, f’, f’’)** are shown here. **a’** displays performance data as in **Fig. 3e**. Brown and gray dashed lines indicate the performance thresholds for upgrading to the next stage and downgrading to the previous stage respectively.

**Supplementary Fig. 3:**
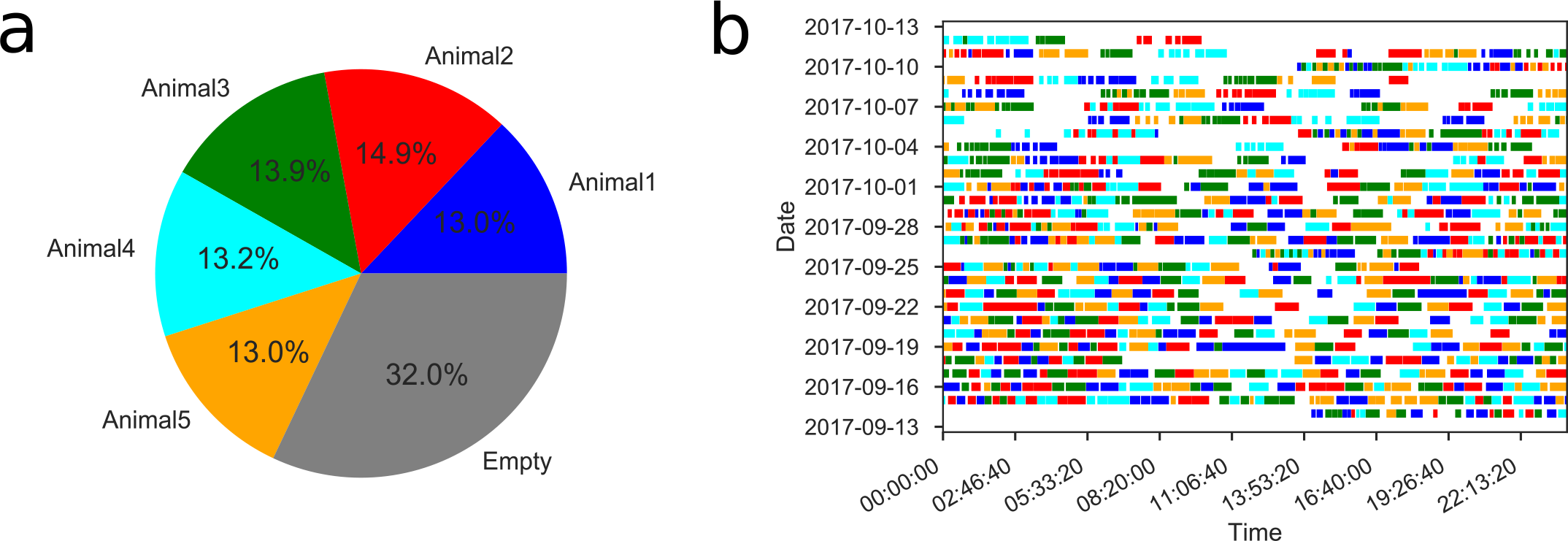
Automated training system allows efficient use of the behavior box. For a sample cohort of five animals, this shows the fraction of time each animal used the behavior box (**a**) and the activity trace of each animal throughout one month.

**Supplementary Fig. 4:**
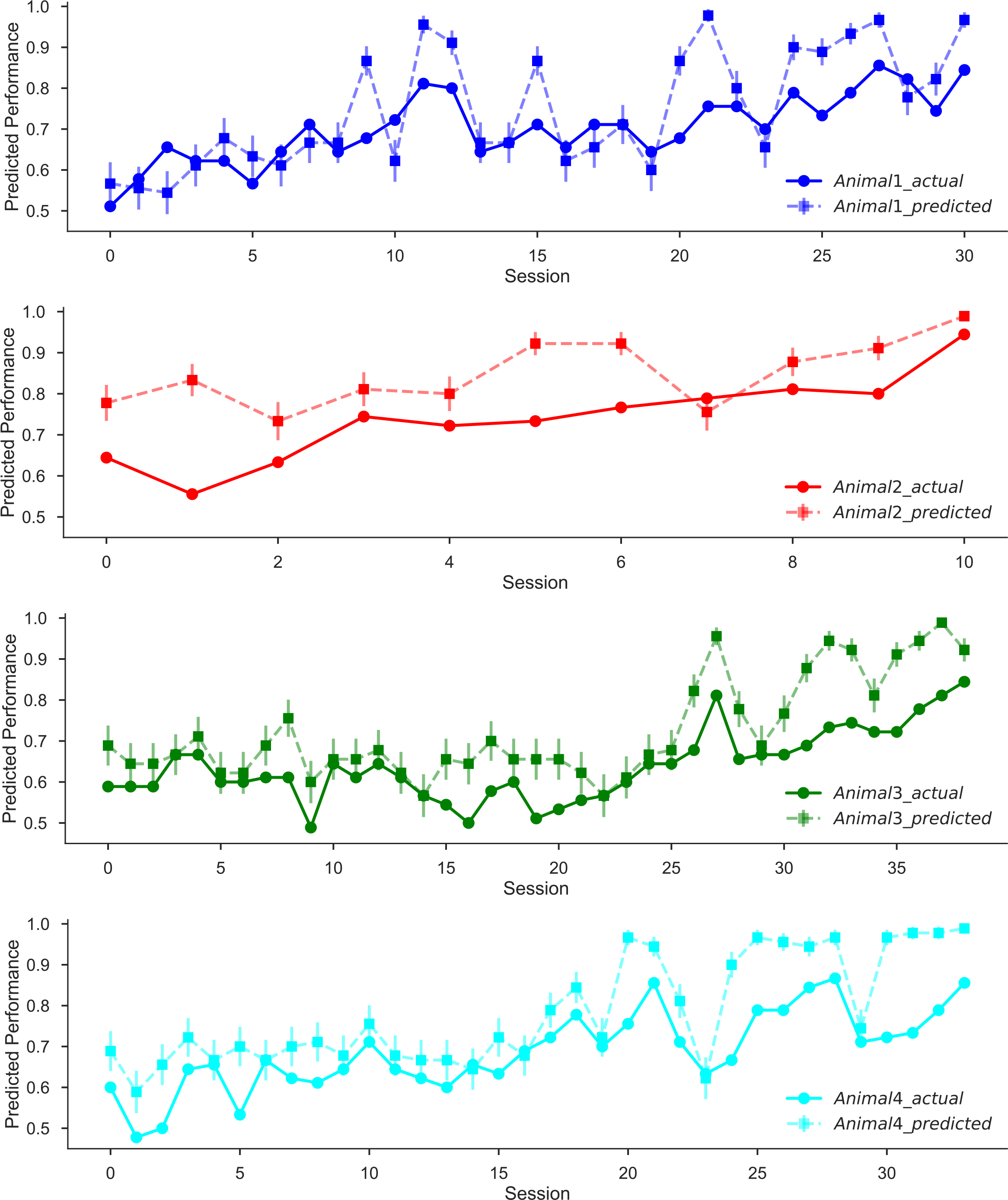
Additional analysis on the iterative generalized linear model’s prediction accuracy. The actual performance and the performance predicted by the model, for each of the four animals. Note that the predictions recapitulate the more prominent fluctuations in the actual learning curves. Error bars indicate standard errors.

**Supplementary Fig. 5:**
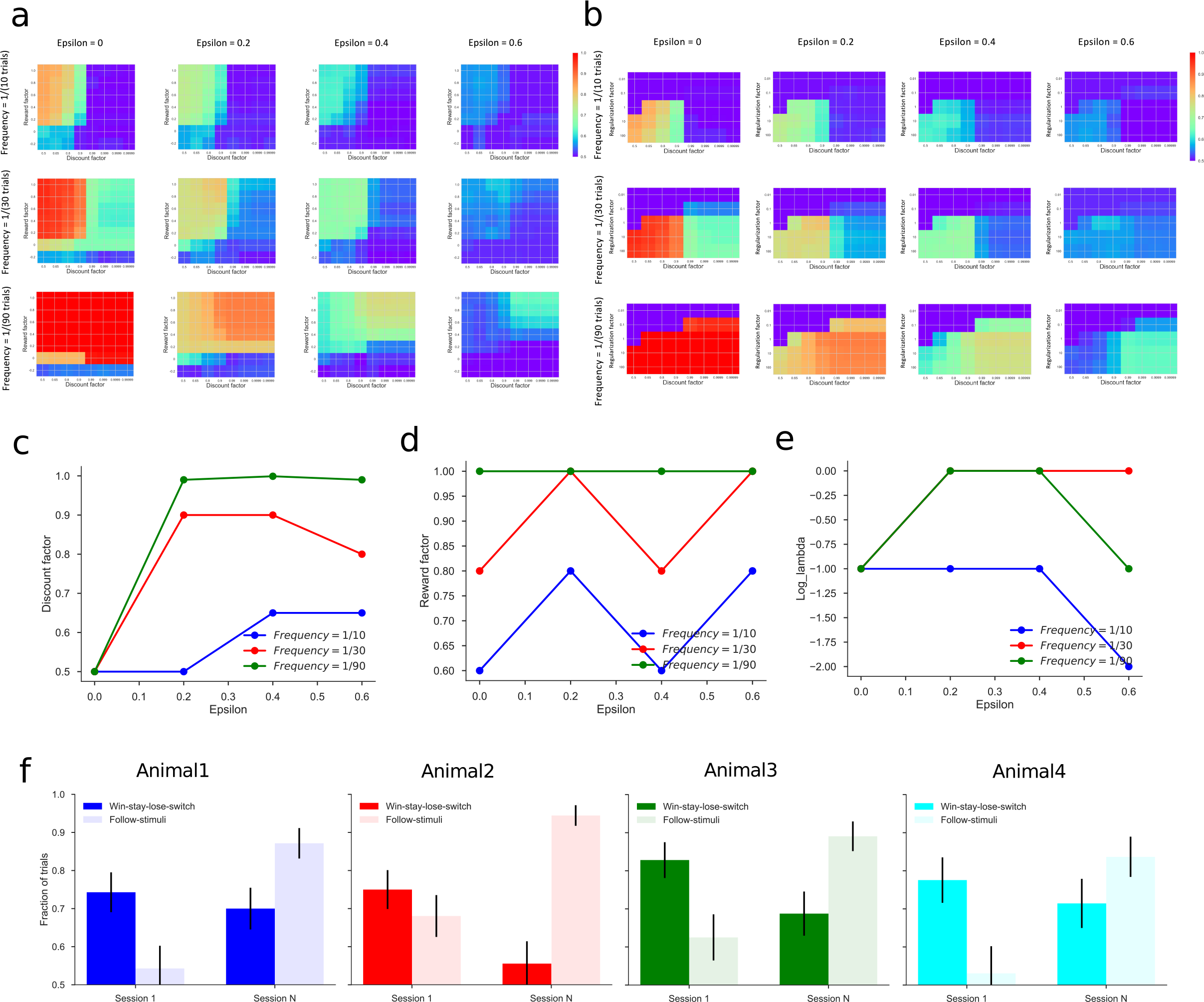
Iterative generalized linear model captures differences between individuals and policy changes. **(a-b)** Hyperparameter selection for the GLMs fitted to the simulated data generated from the ground truth policies. Different values of policy change frequency and noise level (epsilon) lead to different landscapes of the hyperparameters. **(c-e)** Selected temporal discount factor (**c**), reward factor (**d**) and regularization factor (**e**) for different values of policy change frequency and noise level (epsilon). **(f)** Fraction of the trials explained by the two policies (win-stay-lose-switch or WSLS, and following the stimuli) in the first and last sessions, for each of the four animals.

**Supplementary Fig. 6:**
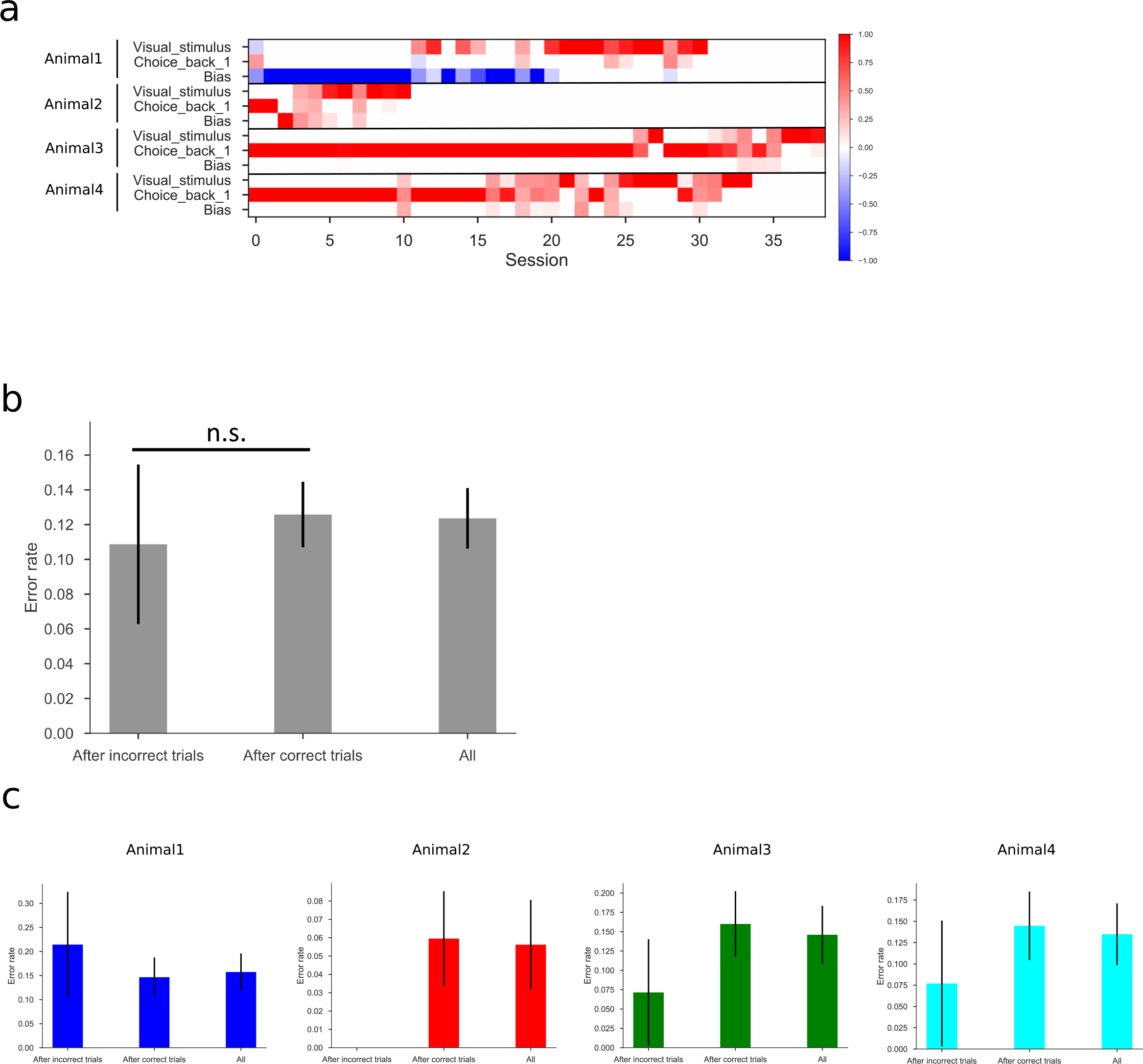
Additional analyses of policy changes during learning. **(a)** Policy matrices over sessions of the four animals. Here the policy matrices are recovered from logistic regression using only the trials following a correct response. Because the reward of the last trial is always +1, the term Reward_back_1 is the same as Bias, and the term RewardxChoice_back_1 is equal to Choice_back_1, so we drop them to avoid redundancy. **(b-c)** Quantification of the error rate during the last session, comparing trials following a correct response to those following a mistake. Averaged over all four animals (**b**) and for each of the four animals (**c**). n.s. indicates not significant.

**Supplementary Fig. 7:**
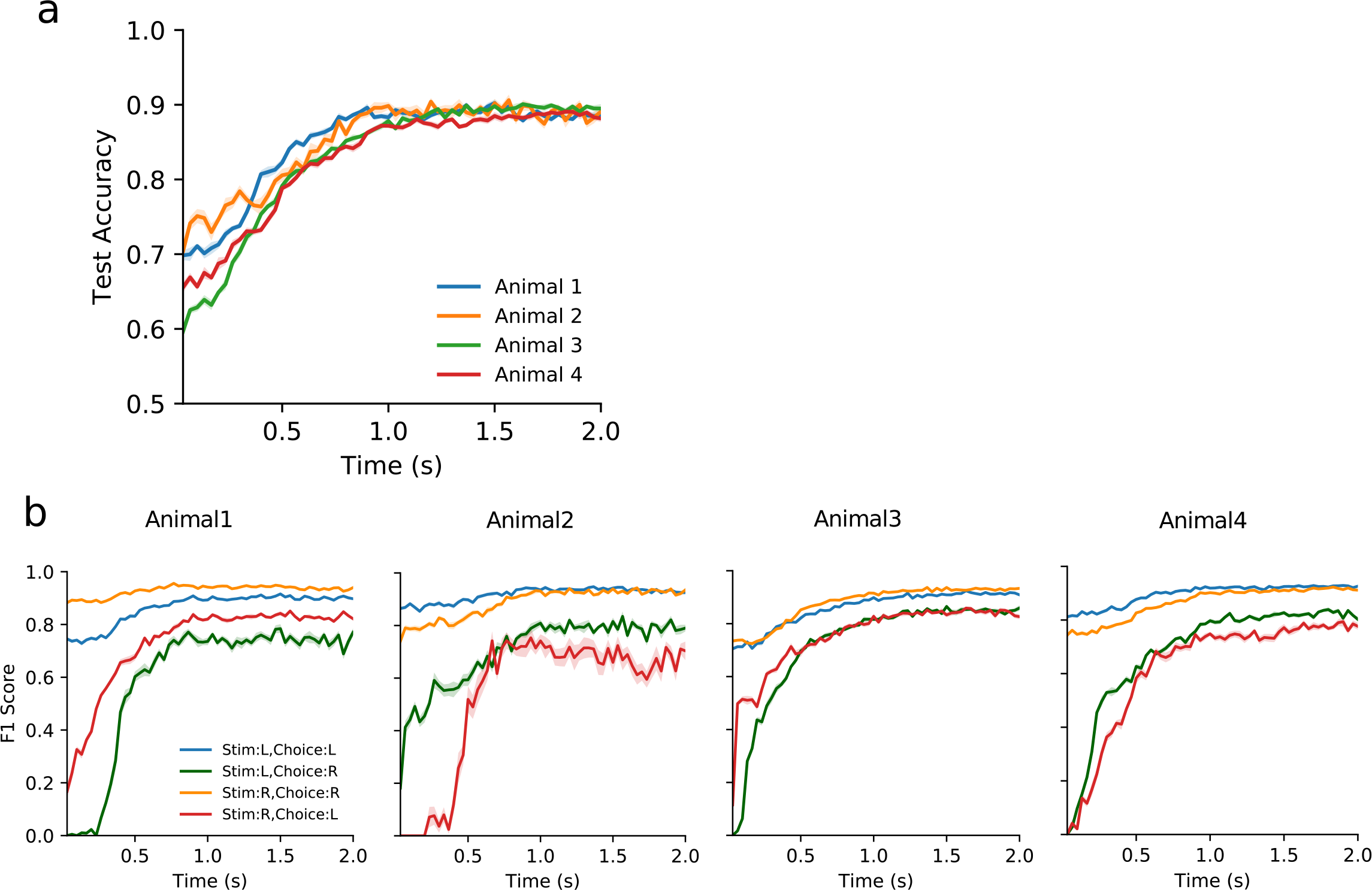
Performance of the support vector machine to infer trial category from mouse trajectories. **(a)** Prediction accuracy of the SVMs for individual animals. **(b)** F1 score of the SVM fitted for the decision categories of each animal. Shaded region denotes standard error. x axis indicates the time starting from when the animal leaves the center port to make a choice. SVMs were trained using features up to a certain time point.

**Supplementary Fig. 8:**
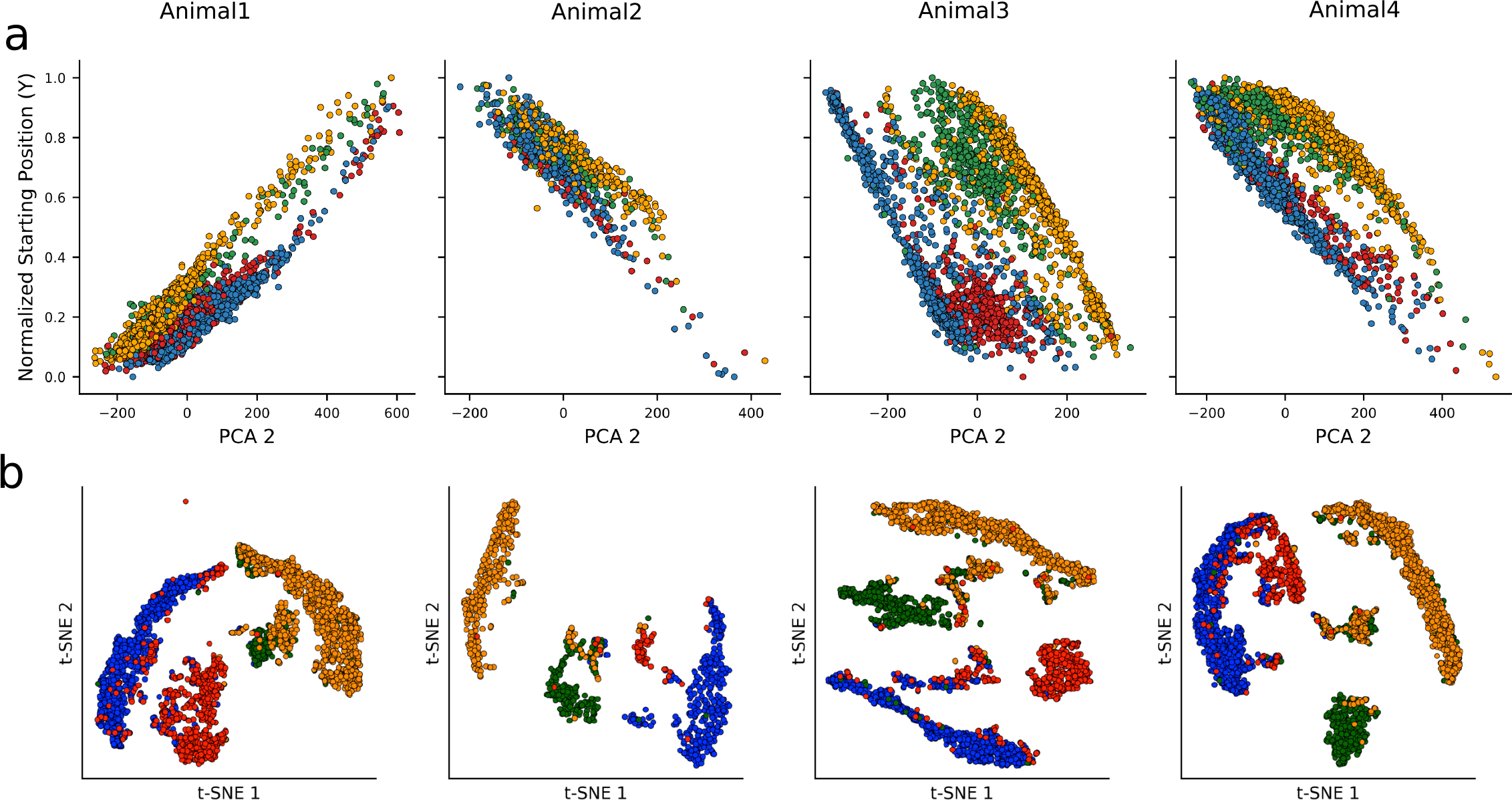
Additional information on the unsupervised analysis of behavior trajectories. **(a)** Scatter plot of starting positions along the left-right axis against PC2 shows correlation between the two. Starting positions are normalized to range from 0 (the leftmost position) and 1 (the rightmost position). **(b)** t-SNE plots with colors indicating different decision categories.

### Supplementary Video 1

https://drive.google.com/open?id=1Ng5s1UhlFRdV4mZ5b1Ot7EUpaEiUXN2_

### Supplementary Video 2

https://drive.google.com/open?id=15qgqM5qOd30kajT-IQkCf8flcF80U2qV

### Supplementary Video 3

https://drive.google.com/open?id=1zqja6_3bA2jO9ap0EWd5t_Z_dVOA8FmA

### Supplementary Video 4

https://drive.google.com/open?id=1wiaaBD-sfZTudDbpRM0ByB2n9bTKyaid

### Supplementary Video 5

https://drive.google.com/open?id=1TQd6N4XW2LDtLbBxkLQezoTJfacNHEZV

